# Cat LCA-*CRX* model, homozygous for an antimorphic mutation has a unique phenotype

**DOI:** 10.1101/2023.03.01.530650

**Authors:** Laurence M. Occelli, Nicholas M. Tran, Shiming Chen, Simon M. Petersen-Jones

**Author notes:** <u>Corresponding author:</u> Simon M. Petersen-Jones, Small Animal Clinical Sciences, Michigan State University, East Lansing, MI 48 824, USA. Molecular and Human Genetics, Baylor College of Medicine, Houston, Tx, USA. **Commercial relationship disclosures:** None.

## Abstract

**PURPOSE:** Human mutations in the *CRX* transcription factor are associated with dominant retinopathies often with more severe macular changes. The *CRX-*mutant cat (*Rdy-A182d2*) is the only animal model with the equivalent of the critical retinal region for high acuity vision, the macula. Heterozygous cats (*CRX^Rdy/+^*) have a severe phenotype modeling Leber congenital amaurosis. This study reports the distinct ocular phenotype of homozygous cats (*CRX^Rdy/Rdy^*).

**METHODS:** Gene expression changes were assessed at both mRNA and protein levels. Changes in globe morphology and retinal structure were analyzed.

**RESULTS:** *CRX^Rdy/Rdy^* cats had high levels of mutant *CRX* mRNA and protein. The expression of photoreceptor target genes was severely impaired while there were variable effects on the expression of other transcription factors. The photoreceptor cells remained immature and failed to elaborate outer segments consistent with the lack of retinal function. The retinal layers displayed a progressive remodeling with cell loss but maintained overall retinal thickness due to gliosis. Rapid photoreceptor loss largely occurred in the macula-equivalent retinal region. The homozygous cats developed markedly increased ocular globe length.

**CONCLUSIONS:** The phenotype of *CRX^Rdy/Rdy^* cats was more severe compared to *CRX^Rdy/+^* cats by several metrics.

**TRANSLATIONAL RELEVANCE:** The *CRX*-mutant cat is the only model for *CRX*-retinopathies with a macula-equivalent region. A prominent feature of the *CRX^Rdy/Rdy^* cat phenotype not detectable in homozygous mouse models, was the rapid degeneration of the macula-equivalent retinal region highlighting the value of this large animal model and its future importance in the testing of translational therapies aiming to restore vision.

## INTRODUCTION

Cone-rod homeobox (CRX) is a transcription factor essential for normal photoreceptor development, function and survival.^1–3^ The N-terminus of the protein contains a homeodomain for DNA binding while the C-terminus contains WSP and OTX-tail domains and controls gene transactivation. CRX directly regulates many genes that are necessary for retinal function, such as those involved in the phototransduction and the visual cycle are directly regulated by CRX. ^4–9^

In humans, *CRX* mutations result in a spectrum of mostly dominant retinopathies with variable severity, ranging from severe childhood-onset forms, such as Leber congenital amaurosis (LCA7), cone–rod dystrophies, retinitis pigmentosa and macular dystrophies.^10–13^ LCA represents approximately 5% of all human inherited retinopathies with a prevalence of 1 in 30,000 to 81,000 newborns.^14, 15^ *CRX* mutations accounts for approximately 2.35% of the cases of LCA.^16^

*CRX* mutations have been categorized into four different classes by the proposed molecular disease mechanism. ^17^ Some mutations, such as those that are null or hypomorphic mutations or those which ablate DNA binding, when present in the heterozygous state result in a mild phenotype (such as a macular dystrophy or mild foveal abnormalities) or no phenotype, while patients that are homozygous for the same mutations have a severe (LCA) phenotype.^18, 19^ Similarly, mice heterozygous for a knockout mutation in *Crx* have a mild phenotype with a slight delay in photoreceptor maturation, whereas mice homozygous for the null mutation have a severe phenotype, equivalent to LCA, with a lack of photoreceptor outer segment development.^20^

The severe dominant LCA phenotypes in patients, and the equivalent in animal models, are associated with expression of a mutant protein with retained DNA binding that has an antimorphic effect.^9, 17^ Class III mutations are antimorphic frameshift/non-sense mutations which escape nonsense mediated decay and produce proteins which have intact DNA binding but lack transactivation activity. In heterozygous animals, this dominant negative CRX protein interferes with the function of the WT allele, and resulting in a more severe dominant phenotype. Three animal models are available for this class of mutations. A knock-in mouse model, *E168d2*, which produces a truncated protein (at 171 of 299 amino acids)^9^, a mouse with a L253X mutation^21^ and the *Rdy* cat, which has a spontaneous frameshift mutation (c.546del, p.Ala185leuTer2) resulting in a truncated protein of 185 amino acids.^22^ The L253X mouse produces a protein with a less truncation of the transactivation domain and a milder phenotype than the other class III models. The *E168d2*/+ mice and *CRX^Rdy/+^* cats have a similar and more severe phenotype associated with over-expression of the mutant allele and higher levels of the mutant than the wild-type protein. ^9, 23^ Development of photoreceptors in *CRX^Rdy/+^* cats is halted resulting in only stunted photoreceptor outer segments. They lack cone electroretinograms (ERG) but initially have delayed and markedly reduced rod-mediated ERGs which are rapidly lost as photoreceptor degeneration progresses. Degeneration is most rapid in the cone-rich *area centralis* which also has a higher packing of photoreceptors than the peripheral retina and is the equivalent of the human macula. The presence of a macula-equivalent retinal region gives the cat model a distinct advantage over the mouse models that do not have similar regional differences in photoreceptor number and distribution.

The purpose of the current study is to report the phenotype of the homozygous (*CRX^Rdy/Rdy^*) cat. As anticipated, the homozygous cat had a more severe decrease in mRNA levels of photoreceptor expressed *CRX* target genes than the heterozygous (*CRX^Rdy/+^*) cat. In contrast to the phenotype of the heterozygote, the homozygotes failed to develop photoreceptor outer segments and had no detectable photoreceptor function. They initially developed relatively normal retinal lamination, although the ONL showed a bilaminar cell body arrangement. With progression they had a loss of photoreceptor nuclei and a marked loss of normal retinal lamination with cell migration and loss and extensive gliosis. The blind homozygous cats developed a marked globe enlargement due to expansion of vitreal cavity also making them a potential model for abnormal globe growth resulting in myopia.

## MATERIALS AND METHODS

### Ethics statement

All procedures were performed in accordance with the ARVO statement for the Use of Animals in Ophthalmic and Vision Research and approved by the Michigan State University Institutional Animal Care and Use Committee.

### Animals

A colony of *CRX^Rdy^* cats maintained at Michigan State University was used for this study and bred to obtain homozygote affected (*CRX^Rdy/Rdy^*), heterozygote affected (*CRX^Rdy/+^*) kittens and wild-type (WT) control cats. Animals were housed under 12D:12L cycles during breeding and 14D:10L the rest of the time. They were fed a commercial feline dry diet (Purina One Smartblend and Purina Kitten Chow; Nestlé Purina, St Louis, MO. USA). Animals studied ranged from 4 weeks to 6 years of age. For n numbers used in the different experiments, please refer to Supplementary Tables S1, S2 and S3.

### Ophthalmic examination and fundus imaging

Full ophthalmic examinations were performed including indirect ophthalmoscopy and wide-field color fundus imaging (Ret-Cam II, Clarity Medical Systems, Inc., Pleasanton, CA, USA). Confocal scanning laser ophthalmoscope (cSLO) fundus imaging (Spectralis OCT+HRA, Heidelberg Engineering Inc., Heidelberg, Germany) was performed under general anesthesia concurrently with spectral domain-optical coherence tomography (SD-OCT) examination (see below).

*In vivo* fluorescein angiography imaged by cSLO was performed in a few animals. Anesthesia, pupil dilation and globe positioning were performed as previously described.^23^ A bolus of 20 mg/kg of 10% sodium fluorescein (Fluorescite 10%, Alcon Laboratories Inc, Fort Worth, Texas, USA) was injected through a 20G catheter in the left cephalic vein followed by a 2mL bolus of Ringer Lactate. Images or video were recorded using a 55° wide-field lens.

### Measurement of globe length

Axial globe length was measured using a combined A- and B-mode ultrasound (A/B Scan System 835, Humphrey, Dublin, CA, USA) under anesthesia. Initially, only the axial globe length was measured, but in later studies the cornea-anterior segment anterior to posterior width, lens width and posterior segment depth were also measured. Measurements in millimeters (mm) were taken from the best A scan and B scan combined on the same images (Fig. 7A).

### Intraocular pressure (IOP)

Intraocular pressure was measured using a TonoVet (Icare Finland Oy, Helsinki, Finland). Measurements were performed at the same time in the morning and 3 recordings averaged for each eye.

### Refraction

Refractive error was assessed using a retinoscope and standard refractive bars.

### Electroretinography (ERG)

Electroretinography was performed on *CRX^Rdy/Rdy^* kittens under general anesthesia as previously described.^23^

### Assessment of retinal morphology and vasculature

#### *In vivo* Spectral Domain-Optical Coherence Tomography

In vivo retinal morphology was assessed by SD-OCT (Spectralis OCT+HRA, Heidelberg Engineering Inc., Heidelberg, Germany) as previously described. ^23^ Total retinal thickness, Receptor+ (REC+; including layers between retinal pigmentary epithelium and outer plexiform layer included)^24^, inner nuclear layer (INL), ganglion cell complex (GCC; including the inner plexiform layer (IPL) and the ganglion cell layer (GCL)) and inner retina (IR; layers between inner nuclear layer and internal limiting membrane) thickness were measured in the center of the *area centralis* and in the four retinal quadrants (at 4 optic nerve head diameter distances from the edge of the optic nerve head superiorly, inferiorly, nasally and temporally) using the Heidelberg Eye Explorer (HEYEX) software.

#### Immunohistochemistry (IHC)

After humane euthanasia, eyes from *CRX^Rdy/Rdy^* and wild-type kittens were collected (at 2, 6, 12 and 20 weeks, and 2 and 3.5 years of age) and processed as previously described.^23^ The antibodies used are listed in Supplementary Table S4.

#### Plastic embedded sections

Eyes were processed for plastic histologic sections and imaged as previously described.^23^ Samples from the dorsal, ventral and *area centralis* regions of the glutaraldehyde fixed eye cup were processed and imaged.

#### Quantitative Reverse Transcriptase-Polymerase Chain Reaction (qRT-PCR)

Retinal samples were collected immediately following euthanasia and globe removal from 2-week-old *CRX^Rdy/Rdy^, CRX^Rdy/+^* and wild-type kittens. Two areas (central and peripheral areas) were dissected as previously described.^23^ Retinal samples were flash frozen and stored at −80°C until RNA extraction, the remaining retina was stored for protein assay. RNA extraction, cDNA synthesis and qRT-PCR reaction were performed as previously described.^9^ RNA quality was assessed, and only samples with an RNA integrity number RIN > 7.0 were used to evaluate gene expression changes. The target genes and primer sequences are shown in Supplementary Table S5.

#### Western blot assay

Processing of retinal samples for Western blot was as previously described.^9, 23^ Monoclonal mouse anti-β-actin antibody (Sigma-Aldrich, Saint Louis, MO, USA) and polyclonal rabbit anti-CRX 119b1 at 1:1000 dilution were used to probe the membranes. Secondary antibodies were goat anti-mouse IRDye 680LT and goat anti-rabbit IRDye 800CW (LI-COR Biosciences, Lincoln, NE, USA) respectively. Fluorescence was detected using the Odyssey Infrared Imager (LI-COR Biosciences, Lincoln, NE, USA). Quantification was performed using Image J (http://imagej.nih.gov/ij/; provided in the public domain by the National Institutes of Health, Bethesda, MD, USA).^25^

#### Statistical analysis

Statistical analysis of IOP, refraction, cDNA level and Western blot fluorescence level data differences were tested for normality (Shapiro-Wilk test for normality). Normally distributed data was analyzed by unpaired 2-tailed Student’s T-test (significance level set at *P* < 0.05), nonparametric data by a Mann-Whitney rank sum test (SigmaPlot 12.0; Systat Software, Inc., San Jose, CA, USA). Student’s T-test was performed when comparing only two groups. A mixed effect model using R studio was used to analyze the data for globe length and SD-OCT measurements; SD-OCT measurements as data was evaluated over time. This was also used to analyze the effect of other factors on IOP and refraction (age) using the equation below.^26^

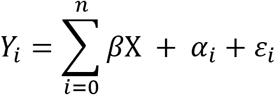

Where *ß* is the parameter vector, X is the independent variable matrix, *α_i_* is the cat level residual, and the *Ɛ_i_* is the individual observation level residual.

## RESULTS

### *CRX^Rdy/Rdy^* retinas overexpressed mutant *CRX* and had a marked reduction in expression of photoreceptor specific genes and variable changes of other retinal transcription factors (Fig. 1)

To investigate molecular changes in the maturing *CRX^Rdy/Rdy^* retina, we measured mRNA levels of *CRX* and other transcription factors involved in photoreceptor development as well as rod and cone specific genes in 2-week-old *CRX^Rdy/Rdy^, CRX^Rdy/+^* and WT kittens (an age at which in wild-type cats photoreceptors are developing inner and outer segments) (Fig. 1. Supplementary Table S6).

**Figure 1.**
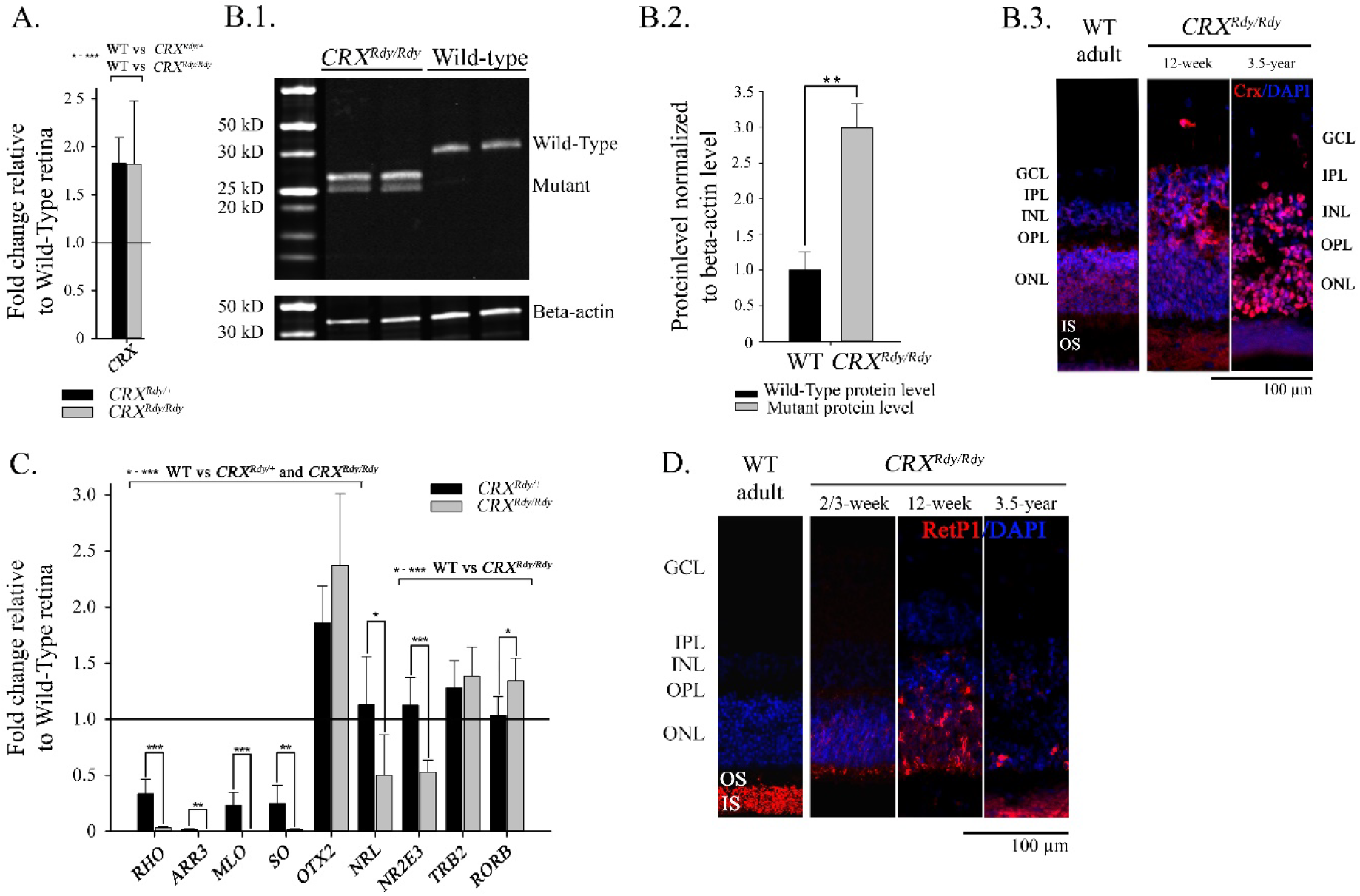
Mutant CRX is overexpressed and alters expression levels of target genes. (**A**) qRT-PCR for total CRX. Overexpression of CRX is apparent in both *CRX^Rdy/+^* and *CRX^Rdy/Rdy^* kittens during retinal maturation (2 weeks of age). Shown relative to WT cat CRX mRNA levels. **(B1)** Western blot for nuclear CRX protein (immunolabeled with antibody 119b1). The truncated mutant CRX is present in the *CRX^Rdy/Rdy^* kitten retina (2 weeks of age). (**B2**) Quantification of total CRX levels. Mutant CRX accumulates to high levels in the *CRX^Rdy/Rdy^* samples compared to the level of WT CRX in the WT control kittens (2 weeks of age). \*\**P* < 0.01. (**B3**) IHC for CRX. Labeling for total CRX shows accumulation in *CRX^Rdy/Rdy^* samples which becomes more pronounced with age. (**c**) Changes in mRNA expression (qRT-PCR) in *CRX^Rdy/Rdy^* retinas compared to *CRX^Rdy/+^* retinas. Expression of photoreceptor genes is further decreased in *CRX^Rdy/Rdy^* retinas compared to *CRX^Rdy/+^* retinas (Rho – rhodopsin; Arr3 cone arrestin; MLO, medium-long wavelength opsin; SO, short wavelength opsin. Expression of transcription factors is also altered: *OTX2* expression is significantly increased compared to WT retinas. *NRL* and *NR2E3* are expressed at lower levels in the *CRX^Rdy/Rdy^* compared to the *CRX^Rdy/+^* and WT kitten, while *TRβ2* and *RORβ* were expressed at higher levels compared to the WT kitten’s retinas. P-values comparing the means *CRX^Rdy/Rdy^, CRX^Rdy/+^* and WT expression levels are:\**P* < 0.05, \*\**P* < 0.01, and \*\*\**P* < 0.001. (**D**) IHC for rhodopsin (RetP1 antibody). In the WT animal this labels the outer segments. In *CRX^Rdy/Rdy^* there is some labeling the very small inner segments (IS) at the 2/3-week time point. There is also some mislocalization with some expression in the ONL cell bodies which was more apparent at 12-weeks. In the adult a few RetP1 positive cells were still apparent.

Levels of total retinal *CRX* mRNA were about 1.8 times higher than WT in both the *CRX^Rdy/Rdy^* and *CRX^Rdy/+^* animals (Fig. 1A). The increased mRNA levels in *CRX^Rdy/Rdy^* animals led to mutant CRX protein levels about 3 times higher than that of the normal CRX in the retinas of WT controls (*P* = 0.005) (Figs. 1B1 and B2). Immunolabeling with an antibody that specifically detects both wild-type and mutant CRX showed that the mutant CRX protein was present across inner and outer nuclear layers in the *CRX^Rdy/Rdy^* cat (shown in Figure 1B3 at 12 weeks of age which is shortly after the age at which the retina is functionally mature in the WT cat). The increased expression of the mutant *CRX* was maintained with age, with strong immunolabelling of retinal cell nuclei across the inner and outer nuclear layres in adult *CRX^Rdy/Rdy^* cats (see representative section at 3.5 years of age in Fig. 1B3). In contrast, the CRX immuno-labeling in the WT adult cat was not so strong and was predominantly detected in the outer nuclear layer.

The *CRX^Rdy/Rdy^* and *CRX^Rdy/+^* mutant cats had marked reduction in mRNA levels of the rod (rhodopsin) and cone (cone arrestin and cone opsins) markers assessed (Fig. 1C). The levels in the *CRX^Rdy/Rdy^* kitten samples were significantly lower than in the *CRX^Rdy/+^* kitten samples. In young animals, despite the marked reduction in rhodopsin mRNA, rhodopsin protein was detected by immunostaining in small protuberances from the outer nuclear layer (ONL) into the subretinal space (Fig. 1D). By 12 weeks, rhodopsin signal expanded and mislocalized to remaining cell bodies in the outer retina (ONL and INL). In adult animals, only a small number of rhodopsin positive cells remained (Fig. 1D). IHC for cone markers (arrestin, and cone opsins) did not label any cells at any ages (Supplementary Fig. S1).

The mRNA levels of CRX-interacting retinal transcription factors (TFs) involved in photoreceptor differentiation were altered in different ways in the *CRX^Rdy/Rdy^* kitten retina compared to WT controls (Fig. 1C): *OTX2, TRβ2* and *RORβ*, which were normally expressed in immature rods/cones were increased, while rod-specific TFs *NRL* and *NR2E3* were decreased.

These results were distinct from those of the *CRX^Rdy/+^* retinas, in which *OTX2* mRNA levels were increased while those for *NRL, NR2E3, TRβ2* and *RORβ* were not significantly different from WT levels. The differences between *CRX^Rdy/Rdy^* and *CRX^Rdy/+^* animals in transcription level changes of all genes examined are shown in Fig. 1C. All *P* values are shown in Supplementary Table S6.

### *CRX^Rdy/Rdy^* retinas developed immature photoreceptors lacking outer segments (Fig. 2) and underwent extensive retinal remodeling (Figs. 3 and 4)

SD-OCT imaging of young *CRX^Rdy/Rdy^* animals showed the development of major retinal layers but the zones such as the ellipsoid zone and interdigitation zone, representing the region of the photoreceptor inner/outer segments and interaction with the retinal pigment epithelium ^27^ could not be detected at any age (Figs. 2A and 3A). Plastic embedded semi-thin sections and TEM sections in young kittens (Fig. 2B and C) showed the presence of small rudimentary inner segments surrounded and in contact with RPE villosities, but no outer segment material was detected. The rudimentary inner segments corresponded to the rhodopsin labelled protuberances from the ONL seen on IHC (Fig. 1D). On SD-OCT, as well as histology and immunohistochemistry sections, the ONL had a bilayered appearance at a young age (4 to 6 weeks). This corresponded to two ONL cell nuclei populations: the outer portion of the ONL had cell bodies with elongated nuclei while the nuclei of cell bodies in the inner portion of the ONL were more circular (Fig. 2B).

**Figure 2.**
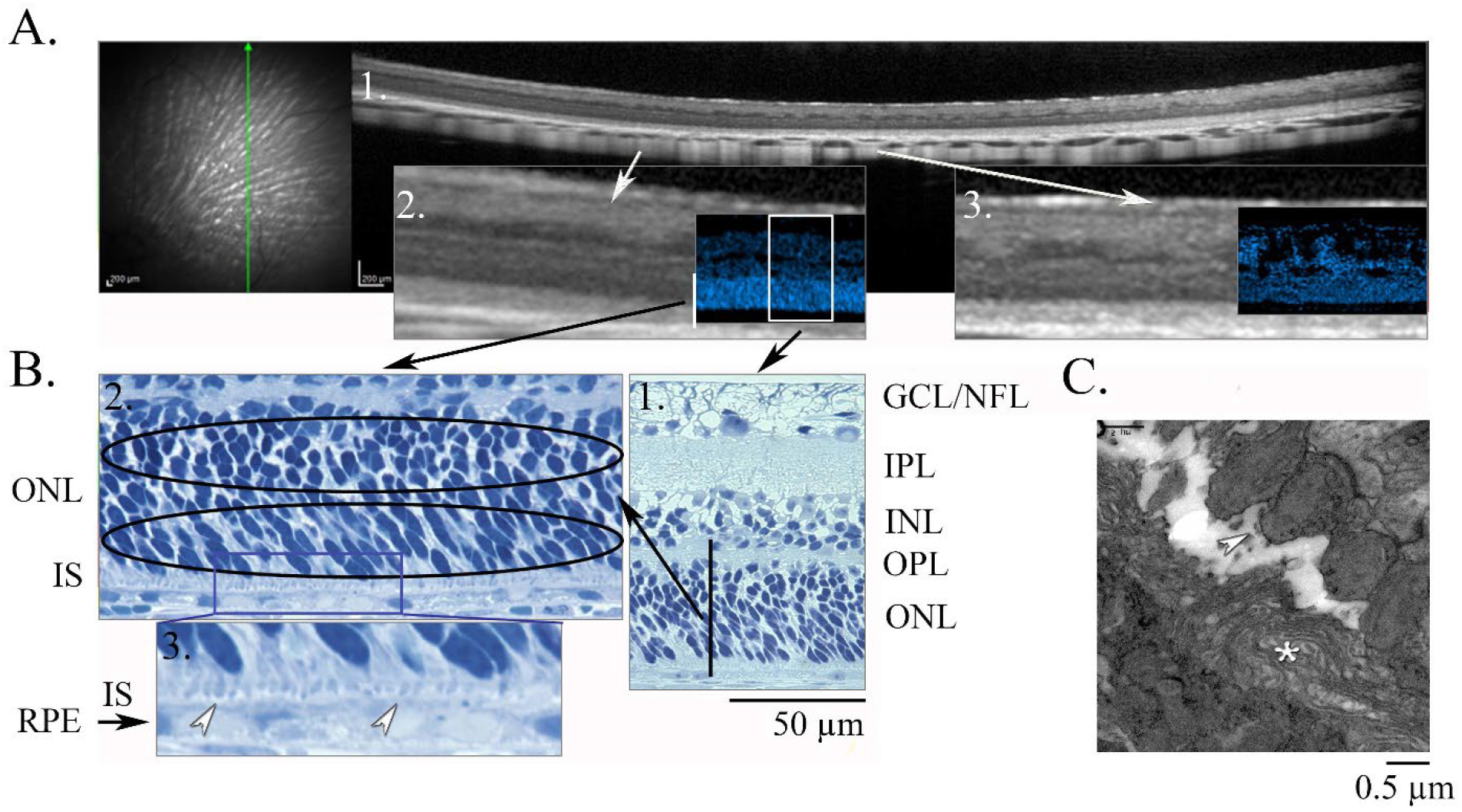
Photoreceptor development is halted in *CRX^Rdy/Rdy^* cats. (**A**) SD-OCT, IHC with DAPI fluorescent staining (**1**) 6-week-old, (**2**) 2-week-old and (**3**) 20-week-old *CRX^Rdy/Rdy^* cats. Relatively normal retinal lamination for the ILM to ONL is apparent in the younger animals. Although the ONL has uneven reflectance. By 20weeks remodeling of retinal layers is apparent on SD-OCT and in the DAPI image. (**B**) Plastic sections at 2 weeks of age. (**1**) Low power view showing relatively normal retinal lamination. (**2**) Magnified view of the outer retina. The ONL nuclei show a bilaminar arrangement. The inner half have nuclei that are round, similar to mature photoreceptors, whereas the outer half has more elongated nuclei similar to immature photoreceptors. (**3**). Higher magnification shows the presence of short vestigial inner segments (white arrowheads) but no obvious outer segment development. (**C**). TEM showing the presence of inner segments (white arrowheads) and RPE villosities (whiteasterisk) but no outer segments.

**Figure 3.**
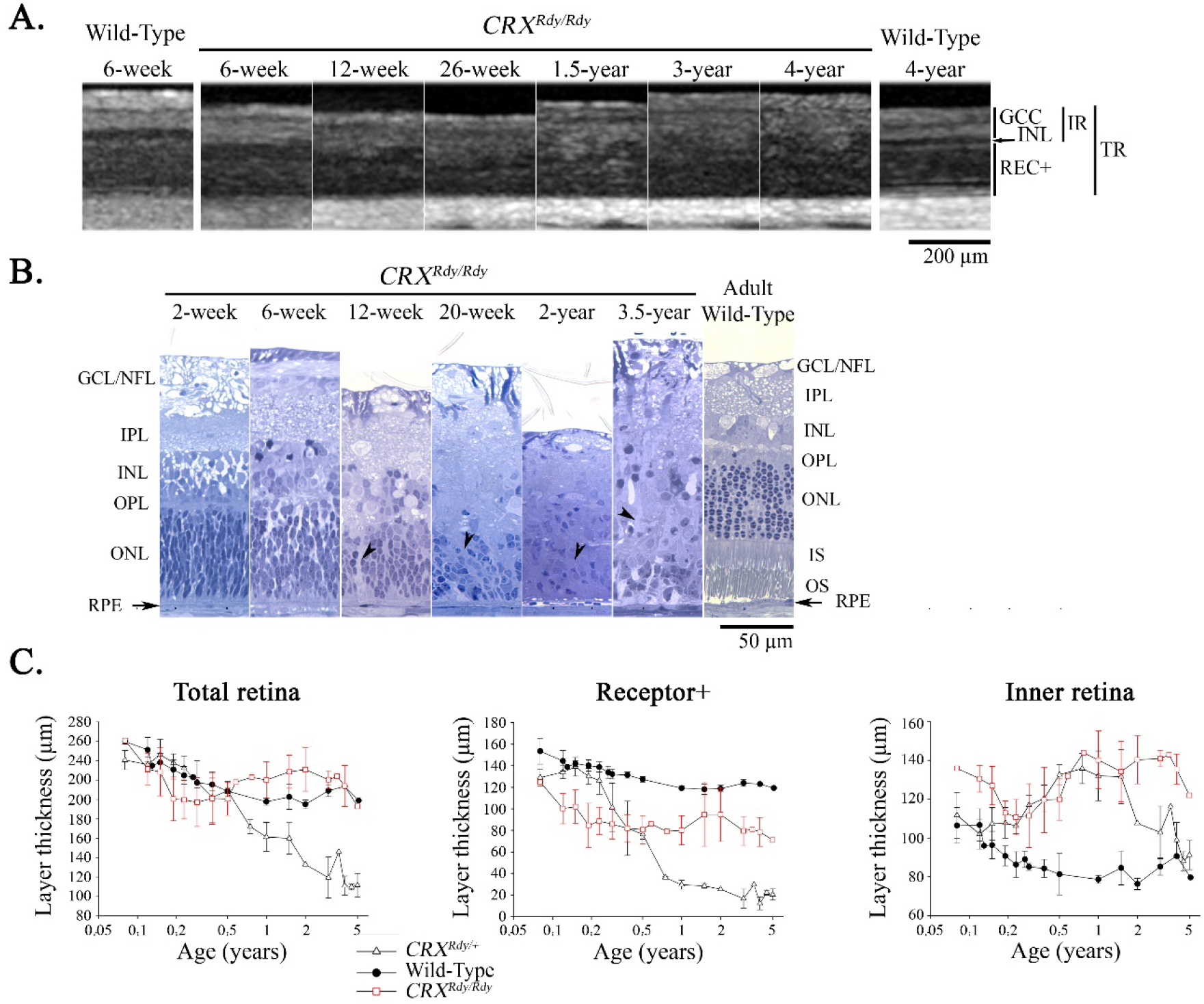
Extensive remodeling of *CRX^Rdy/Rdy^* retina occurs with age. (**A**). SD-OCT comparing dorsal region imaging of retina of *CRX^Rdy/Rdy^* cats with WT controls. At 6-weeks the SD-OCT of the *CRX^Rdy/Rdy^* cat and WT are similar except for the absence of zones representing the outer segments. With age the definition of layers is lost in the *CRX^Rdy/Rdy^* cat although overall retinal thickness is maintained and a demarcation between inner and outer retina can still be discerned. (**B**) Plastic sections showing the loss of cell bodies and loss of retinal lamination with age. Overall retinal thickness is maintained most likely due to glial activation (black arrowheads indicate glial extensions between remaining cell bodies – also see Figure 4B). (**C**) Thicknesses of retinal layers in the dorsal region from SD-OCT imaging from *CRX^Rdy/Rdy^, CRX^Rdy/+^* and WT cats (mean +/- SD). Note the maintenance of the total retinal thickness but thinning of REC+ and thickening of the IR in the *CRX^Rdy/Rdy^* cat. In the *CRX^Rdy/+^* cat progressive outer retinal thinning occurs – as shown on REC+ graph. **Key:** TR, Total retina; REC+; Receptor+; IR, Inner retina; INL, Inner retina layer; GCC, Ganglion cell complex. GCL/NFL, ganglion cell layer/nerve fiber layer; IPL, inner plexiform layer; INL, inner nuclear layer; OPL, outer plexiform layer; ONL, outer nuclear layer; OS, outer segment; IS, inner segment; RPE, retinal pigment epithelium.

The normal lamination of the retina became less apparent as early as 12 weeks of age and further deteriorated with age, although the overall retinal thickness was maintained. Figure 3 illustrates the changes in SD-OCT appearance and retinal morphology of the dorsal central retinal region with age. Changes in thicknesses in the other 3 quadrants examined showed a similar pattern (data not shown). The *area centralis* showed a different pattern of retinal layer thickness changes and is considered separately below. The maintenance of overall retinal thickness with age was different from the findings in the *CRX^Rdy/+^* cats which had a progressive retinal thinning predominantly driven by more severe outer retinal thinning (Fig. 3C). Both *CRX^Rdy/+^* and *CRX^Rdy/Rdy^* cats showed noticeable thickening of the inner retina with age. In the *CRX^Rdy/Rdy^* cats this compensated for the outer retinal thinning meaning that the overall retinal thickness remained close to that of WT cats.

The plastic sections in Figure 3B show the loss of retinal cells in the ONL and INL and the apparent appearance of glial cell processes extending to within the ONL (Fig. 3B, black arrowheads). IHC (Fig. 4B) showed marked GFAP immunolabelling by 12 weeks of age. In the more mature animals GFAP signal was even more extensive throughout the retina showing extensive Müller cell activation and retinal gliosis.^28–30^

**Figure 4.**
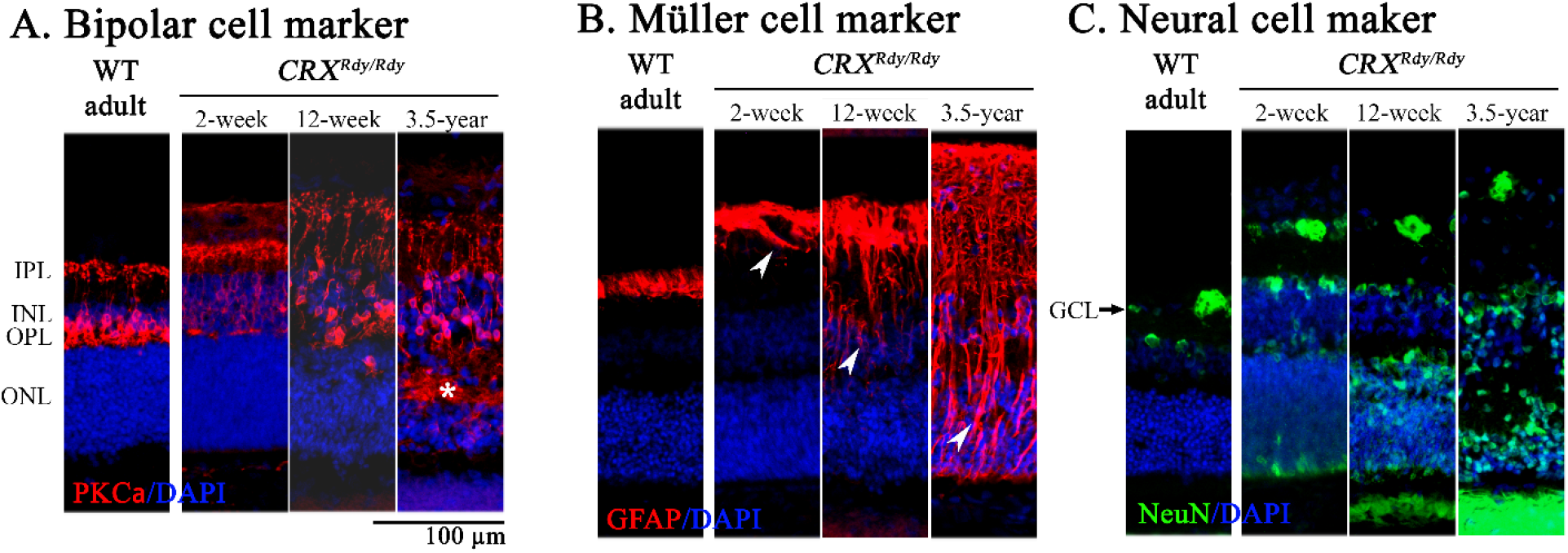
IHC showing inner retinal cell changes with time in *CRX^Rdy/Rdy^* cat. (**A**). PKCalpha which labels bipolar cells shows that with age there is disorganization of the labeled cells with dendrites invading the ONL and forming a matrix within it (white star). (**B**). GFAP is a marker of Muller glia activation. The normal cat has labeling in the region of the ganglion cell layer. With age the is a progression of GFAP positive processes throughout the retina (arrowheads show Muller cell processes which with age tend to replace other cell type. **(C)** Immunolabeling with NeuN antibody showed apparently normal labeling of ganglion cells and some INL cells as in the WT retina, but there was also some abnormal labeling through the ONL nuclei. OS, Photoreceptor outer segment; IS, Photoreceptor inner segment; ONL, Outer nuclear layer; OPL, Outer plexiform layer; INL, Inner nuclear layer; IPL, Inner plexiform layer; GCL, ganglion cell layer.

Immunolabeling of bipolar cells showed extensive migration and branching of cells with age. By 3.5 years of age PKC alpha positive bipolar cells extended throughout the remaining ONL (Fig. 4A). NeuN, a neuronal marker, labeled ganglion cells as in normal retina but it also abnormally labeled inner retinal cells and ONL nuclei at all age tested in the *CRX^Rdy/Rdy^* retina (Fig. 4C).

### *CRX^Rdy/Rdy^* retinas showed early degeneration of photoreceptors in the *area centralis* (Fig. 5)

SD-OCT across the area centralis showed an early thinning of the ONL (Fig. 5A and B). This developed at an earlier age than seen in the *CRX^Rdy/+^* kittens (Fig. 5A). The SD-OCT finding was confirmed by histology (Fig 5. B lower histology images). Supplementary Figure S2 shows heat maps for retinal layer thickness of the area centralis from SD-OCT measure showing the progressive thinning of the outer retina while inner retina thickness increases.

**Figure 5.**
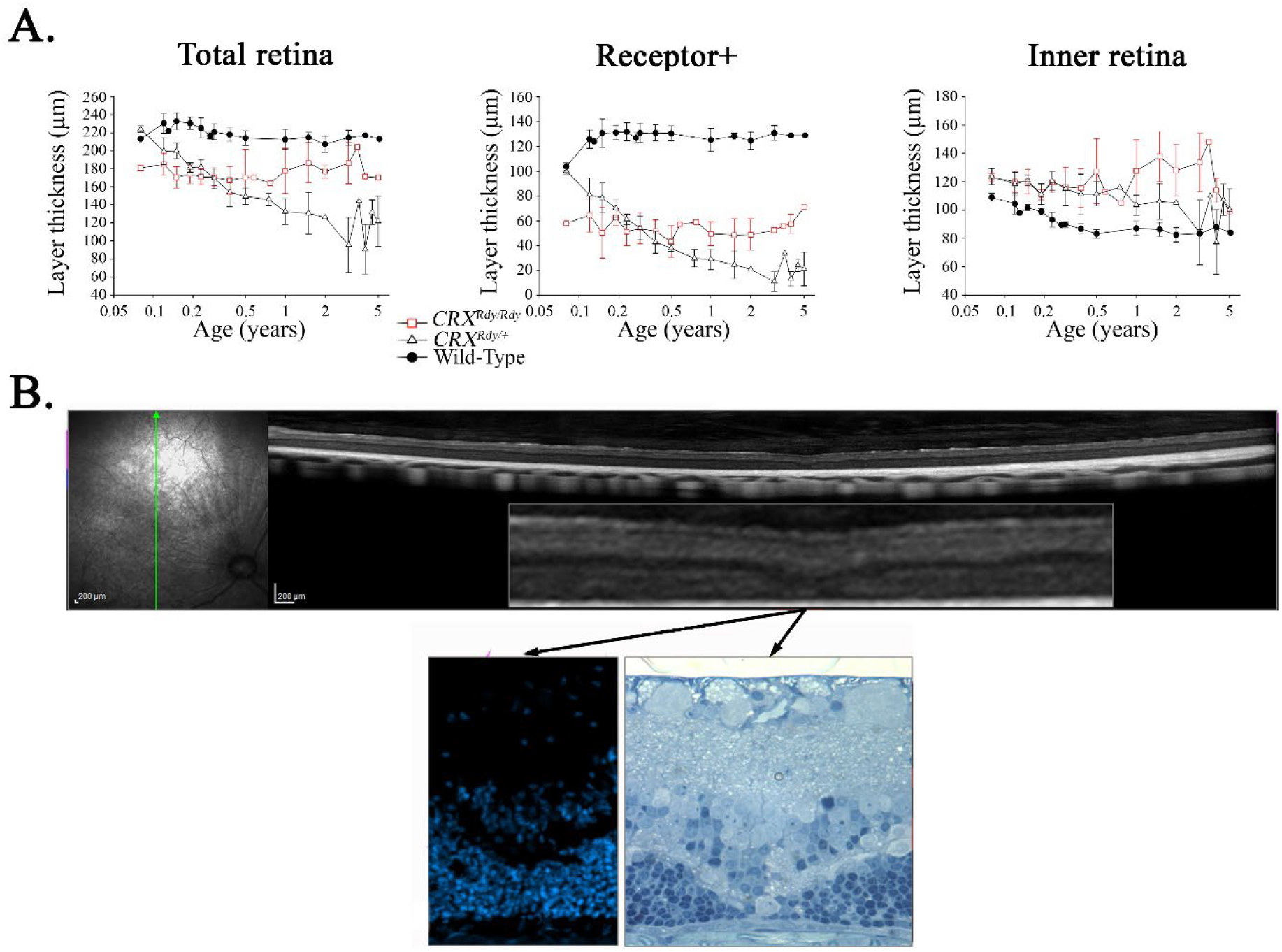
Early degeneration in the center of the area centralis. (**A**). SD-OCT layer measurement in the center of the area centralis from *CRX^Rdy/Rdy^, CRX^Rdy/+^* and WT cats (mean +/- SD). Note the early and marked loss of REC+ in the *CRX^Rdy/Rdy^* cat wile inner retina thickens and total retinal thickness is maintained. (**B**). SD-OCT scan vertically through the area centralis of a 20-week-old *CRX^Rdy/Rdy^*, cat. The insert shows a magnified image of the area centralis with outer retinal thinning. Below a DAPI labeled frozen section and plastic section are shown. These show the small region of marked ONL thinning present at the center of the *area centralis*.

### *CRX^Rdy/Rdy^* kittens lack retinal function

The *CRX^Rdy/Rdy^* cats were blind. They had no dazzle reflex at any age, lacked visual tracking responses and did not develop a menace response. Dark- and light-adapted ERGs assessed at multiple time points from 4 to 20 weeks of age failed to elicit any measurable response (data not shown). A slow and reduced pupillary light reflex was initially present. Nystagmus was not noted at any age.

### *CRX^Rdy/Rdy^* cats developed tapetal hyperreflectivity and local choroidal atrophy but maintained superficial retinal vasculature (Fig. 6)

Color fundus images are shown in Figure 6. The important features are that hyperreflectivity of the tapetal fundus became apparent with age (Fig. 6A). This could be discerned as early as 12 weeks of age (Fig. 6A) and in fact the appearance of the tapetum was never normal having a generalized abnormal “sheen” when compared to WT cats. Tapetal hyperreflectivity indicates that there is less attenuation of light as it passes through the retinal layers and is reflected back from the tapetum. With progression some loss of tapetal tissue became apparent in a pattern radiating out from the optic nerve head; compare the fundus images in Figure 6A at 26 weeks and 3 years of age which are from the same animal and show the progression of these lesions. With the loss of tapetum, choroidal vessels could be visualized. SD-OCT imaging showed tapetal and choroidal thinning in those areas (data not shown).

**Figure 6.**
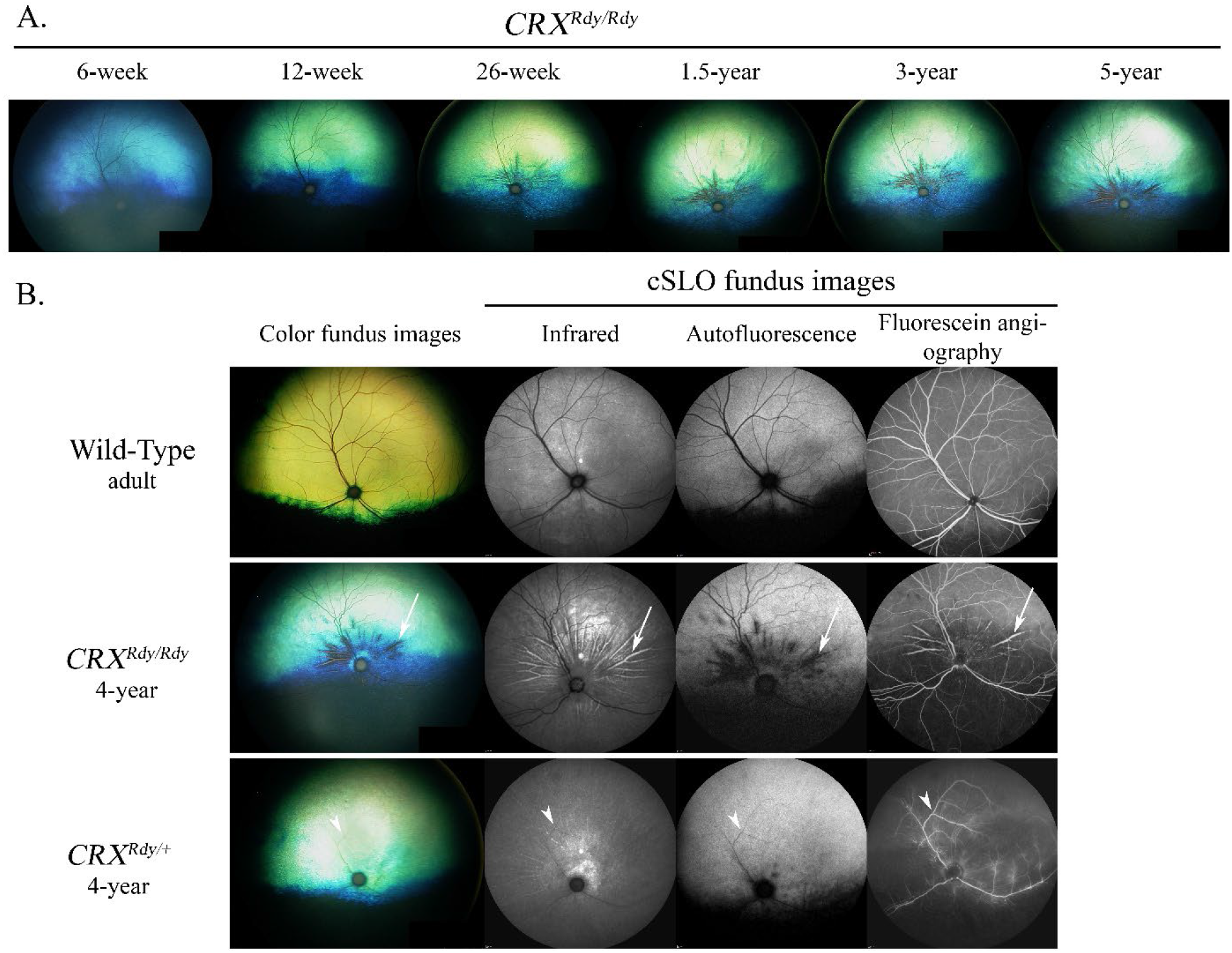
*CRX^Rdy/Rdy^* cats have good preservation of superficial retinal vasculature compared to the *CRX^Rdy/+^* cats. (**A**) Fundus images of *CRX^Rdy/Rdy^* showing development of tapetal hyperreflectivity with age which usually indicates retinal thinning. There is also tapetal thinning close to the optic nerve head allowing tapetal vasculature to be visualized. (**B**) comparison of WT cat with *CRX^Rdy/Rdy^* and *CRX^Rdy/+^* 4-year-old cats. The color fundus images are followed by cSLO infrared, autofluorescence and fluorescein angiography images. The *CRX^Rdy/Rdy^* cat shows a lack of tapetum as dark streaks radiating from the optic nerve head. The white arrow indicated a choroidal vessel that is exposed. Compare the vasculature to that of the *CRX^Rdy/+^* cat which has only the major vessels still detectable. The white arrowhead indicates the same vessel for the three types of imaging.

The superficial retinal vasculature was well maintained in the *CRX^Rdy/Rdy^* cats. Figure 6B compares color fundus images and cSLO and fluorescein angiography images between a representative adult WT cat and age matched *CRX^Rdy/Rdy^* and *CRX^Rdy/+^* cats. The *CRX^Rdy/Rdy^* cat has developed less hyperreflectivity of the tapetum (color images) than the *CRX^Rdy/+^* cat, and shows better preservation of superficial retinal vasculature. One region of tapetal loss in the *CRX^Rdy/Rdy^* cat is indicated with a white arrow and fluorescein angiography highlights the exposed underlying choroidal vessel. The relative preservation of superficial retinal vessels in the *CRX^Rdy/Rdy^* cats compared to *CRX^Rdy/+^* cats is clearly illustrated in the fluorescein angiography panel. The *CRX^Rdy/+^* cat had markedly attenuated retinal vasculature with mild leakage of fluorescein from remaining superficial retinal vessels.

### *CRX^Rdy/Rdy^* cats develop an increased globe length and a myopic refractive error (Fig. 7)

*CRX^Rdy/Rdy^* cats had a significantly increased axial globe length compared to *CRX^Rdy/+^* and control WT cats (Fig. 7) (*P* < 0.002 and = 0.003, respectively). For example, at one year of year the *CRX^Rdy/Rdy^, CRX^Rdy/+^* and WT cats had an axial globe length of respectively 23.1 ± 0.4, 19.5 ± 0.3 and 20.1 ± 0.4 mm. This difference in globe size was obvious on ocular ultrasound (Fig. 7A, bottom panel), and in the enucleated eye (Figs. 7B). Scatter plots in Figures 7C and D show the changes with age. The increase in axial globe length was due to an increase in the posterior segment length (*P* < 0.0001) (Fig. 7D). The anterior-posterior depth of the anterior chamber and the width of the lens did not differ between genotypes (data not shown). A difference in axial globe length was also present between *CRX^Rdy/+^* and WT cats (*P* = 0.02), with WT having a slightly greater axial length. It should be noted that the *CRX^Rdy/Rdy^* group included 55 males (M) and 13 females (F) (each time point considered independently), the WT group included 94 M and 54 F and the *CRX^Rdy/+^* group in 44 M and 106 F. It is possible that the difference between the *CRX^Rdy/+^* and the WT group was due to high number of females that were of smaller size compared to the males. For example, at 3 years of age female and male *CRX^Rdy/+^* cats had a mean axial globe length of 20.4 ± 0.13 and 21.50 ± 0.34 mm respectively, a difference which was statistically significant (*P* ≤ 0.001).

**Figure 7.**
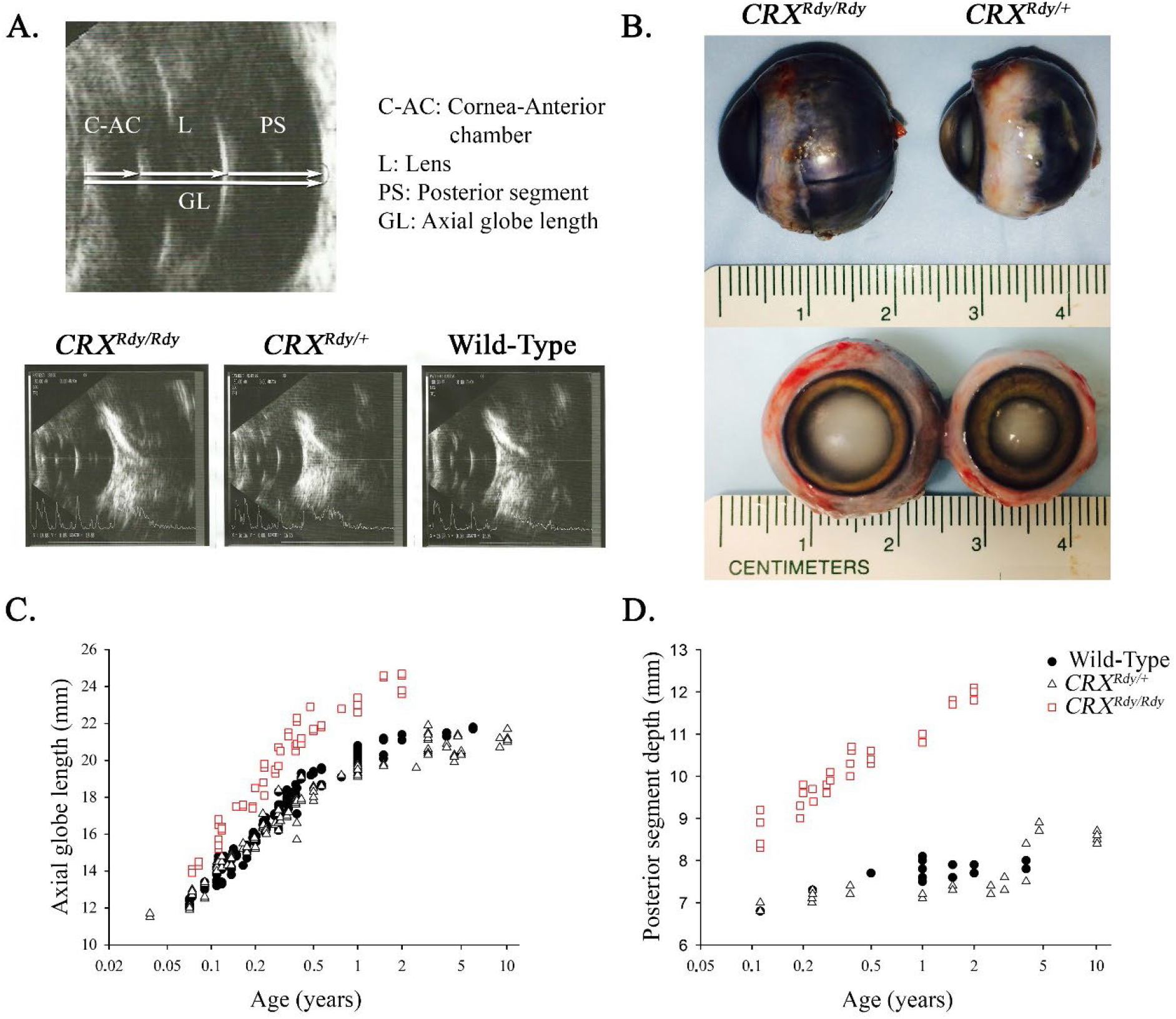
*CRX^Rdy/Rdy^* cats develop globe enlargement. (**A**) Ultrasound images showing the measurements performed and below representative images from the different genotypes (each 12 weeks of age). (**B**) Gross pictures of enucleated globes from age-matched (12 weeks of age) animals showing the much larger globe in the *CRX^Rdy/Rdy^* compared to *CRX^Rdy/+^* cat. (**C**) Scatter plots of the axial length of the 3 genotypes with age. (**D**) Scatter plots of the posterior segment length of the 3 genotypes with age.

There was no difference in intraocular pressures (IOPs) between *CRX^Rdy/Rdy^* and WT cats (16.0 ± 1.8 and 15.4 ± 1.6 mmHg, respectively *P* = 0.577). The change in axial globe length did alter the refractive state of the eye. The *CRX^Rdy/Rdy^* cats had a refractive error of a mean of −14 ± 1.1 D compared to −0.9 ± 0.4 D in the WT cats (*P* = 0.004). A refractive error was also present in the *CRX^Rdy/+^* cats with an average of −2.7 ± 0.8 D (*P* < 0.001 compared to *CRX^Rdy/Rdy^* cats and *P* = 0.002 compared to WT cats). When analyzed with a mixed effect model, age had a significant influence in the difference between the *CRX^Rdy/+^* cats and both the *CRX^Rdy/Rdy^* and WT cats (*P* = 0.003). The eye (left or right) did not have an effect on the difference between groups nor did it have an effect on refraction or IOP.

## DISCUSSION

This study adds to previous studies characterizing the *Rdy* cat. While phenotypic descriptions of the heterozygous *CRX^Rdy/+^* cat have been previously reported, there had been no studies reporting on the homozygous *CRX^Rdy/Rdy^* cat.^23, 31–34^ In keeping with previous studies of Class III *Crx* mouse mutant models, the *CRX_Rdy/Rdy_* cat showed an over-expression of the mutant CRX transcript and protein that we predict to exert an antimorphic effect on developing photoreceptor cells. The *Rdy* feline model offers important advantages due to the presence of a macula-like retinal region allowing for further characterization of the phenotype of Class III *CRX* mutations.

### Molecular mechanism underlying *CRX^Rdy/Rdy^* phenotype

The Rdy mutation introduces a premature stop codon ^22^ in *CRX* with the mutant transcript escaping nonsense-mediated decay and resulting in overexpression of the mutant allele (Fig. 1A and B).^9, 35, 36^

Lack of normal CRX activity resulted in a marked decrease in cone and rod transcripts in the *CRX^Rdy/Rdy^* cat retinas (Figs. 1C and D). It also had effects on the levels of other transcription factors involved in photoreceptor development. *CRX, OTX2* and *RORβ* showed an increase in expression. The increase in *OTX2* expression may be due to a retroactive feedback mechanism compensating for the lack of other transcription factors. *NRL* and *NR2E3* expression level was decreased, perhaps contributing to the partial failure in maturation of the photoreceptor nuclei. Although the cone maturation factor *TRβ2*, is still expressed, there was a lack of cone opsin expression and progressive cone degeneration, suggesting CRX is required for cone maturation and maintenance. While Rdy clearly inhibit the expression of many rod and cone-specific genes necessary for vision, photoreceptor transcription factors showed both positive and negative regulation, as these transcriptional network interactions are complex, this regulatory process is still not completely understood.^2, 37^

### The *CRX^Rdy/Rdy^* kitten is a model for severe *CRX*-LCA retinopathies

The *CRX^Rdy/Rdy^* cat phenotype is characterized by blindness from birth, with a sluggish PLR, and no dazzle reflex or menace response. We suspected that the slow residual PLR was driven by melanopsin containing ganglion cells.^38^ Ophthalmoscopic features of the model included the development of tapetal hyperreflectivity, which is considered an indicator of generalized retinal thinning. However, there was not a significant decrease in total retinal thickness so the tapetal hyperreflectivity likely either resulted from the almost total lack of cone and rod opsins that in a normal eye would absorb photons passing through the retina or the extensive cellular degeneration and remodeling that occurred. The photoreceptors of the *CRX^Rdy/Rdy^* cat only developed very short rhodopsin positive inner segments (Fig. 1D) before photoreceptor maturation became halted. The photoreceptor nuclei consisted of two populations with different nuclear morphology: one which population had oval-shaped nuclei comparable to the appearance in an immature retina while the other showed a more mature morphology (round nuclei). It appeared that the normal migration of cell bodies of photoreceptors that occurs during maturation of the retina did not occur. The lack of photoreceptor maturation was reflected in the complete lack of detectable ERG responses. Interestingly, photoreceptor degeneration was not as rapid as in the *CRX^Rdy/+^* cat, although degeneration and marked cell migration occurred. There was initially relatively normal stratification of the retinal layers, but over time there was extensive activation of Müller cells and sprouting and migration of bipolar cells (Figs. 3 and 4). This retinal remodeling resulted in some thickening of the inner retinal layers, which counteracted the thinning of the outer retain and thus preserved the overall retinal thickness (Fig. 3).^28–30, 39^ This retinal remodeling was more severe than in the *CRX^Rdy/+^* feline phenotype ^23^ but was similar to the *E168d2/d2* homozygous mouse.^17^ The relative lack of overall retinal thinning may account for the striking persistence of retinal vasculature until advanced disease stages (Fig. 6). The *area centralis* had slightly different changes compared to the peripheral retina with thinning of the total retina and receptor+ layers (Fig. 5 and Supplementary Fig. S2) with early severe loss of photoreceptor nuclei in the center of the *area centralis*.

The phenotype of the *CRX^Rdy/Rdy^* cat is quite different to that of the heterozygote, not being simply a more severe version. The absence of photoreceptor OS development might be responsible for the relatively slow photoreceptor nuclei loss compared to the *CRX^Rdy/+^* cat. In the *CRX^Rdy/+^* cats, the degeneration of photoreceptors expressing phototransduction proteins and retinal degeneration could result from altered phototransduction cascade dynamics and mislocalization of opsins.^40–45^

### The *CRX^Rdy/Rdy^* kitten develops significant globe enlargement without glaucoma

The *CRX^Rdy/Rdy^* cat has a significant increase in axial globe length with resulting myopia due to an increase in posterior segment length (Fig. 7).^46–49^ Abnormal globe length is a well-recognized feature in animals with abnormal visual input.^50, 51^ The increase in globe size may also contribute to the choroidal thinning that was noted in the homozygous cats (Fig. 6).^52^

The *CRX^Rdy/Rdy^* cat provides a large animal model for investigating scleral growth factors implicated in myopia development and for the severe dominant *CRX* mutations associated with over expression of a mutant transcript with an antimorphic effect.

## ACKNOWLEDGEMENTS

The authors would like to thank Dr. Cheryl Craft for the donation of the hCAR antibody, Dr. Alicia Withrow for her help with semi-thin sections, Dr. Wenjuan Ma for statistical advice and the MSU CVM RATTS group for their help with animal handling.

Supported by National Institutes of Health Grants EY012543 and EY025272-01A1 (SC), EY002687 (P30 Core Grant) (Washington University Department of Ophthalmology and Visual Sciences (WU-DOVS)), EY013360 (T32 Predoctoral Training Grant) (WU), unrestricted funds from Research to Prevent Blindness (WUDOVS), Foundation Fighting Blindness (SC), Hope for Vision (SC), George H. Bird and “Casper” Endowment for Feline Initiatives (LMO and SMPJ), Michigan State University Center for Feline Health and Well-Being (LMO and SMPJ), and Myers-Dunlap Endowment (SMPJ).

## REFERENCES

1. Furukawa T, Morrow EM, Cepko CL. Crx, a novel otx-like homeobox gene, shows photoreceptor-specific expression and regulates photoreceptor differentiation. Cell 1997;91:531–541.

2. Hennig AK, Peng GH, Chen S. Regulation of photoreceptor gene expression by Crx-associated transcription factor network. Brain Res 2008;1192:114–133.

3. Morrow EM, Furukawa T, Raviola E, Cepko CL. Synaptogenesis and outer segment formation are perturbed in the neural retina of Crx mutant mice. BMC neuroscience 2005;6:5.

4. Chau KY, Chen S, Zack DJ, Ono SJ. Functional domains of the cone-rod homeobox (CRX) transcription factor. J Biol Chem 2000;275:37264–37270.

5. Chen S, Wang QL, Nie Z, et al. Crx, a novel Otx-like paired-homeodomain protein, binds to and transactivates photoreceptor cell-specific genes. Neuron 1997;19:1017–1030.

6. Peng GH, Chen S. Crx activates opsin transcription by recruiting HAT-containing co-activators and promoting histone acetylation. Hum Mol Genet 2007;16:2433–2452.

7. Corbo JC, Lawrence KA, Karlstetter M, et al. CRX ChIP-seq reveals the cis-regulatory architecture of mouse photoreceptors. Genome Res 2010;20:1512–1525.

8. Livesey FJ, Furukawa T, Steffen MA, Church GM, Cepko CL. Microarray analysis of the transcriptional network controlled by the photoreceptor homeobox gene Crx. Curr Biol 2000;10:301–310.

9. Tran NM, Zhang A, Zhang X, Huecker JB, Hennig AK, Chen S. Mechanistically Distinct Mouse Models for CRX-Associated Retinopathy. PLoS Genet 2014;10:e1004111.

10. Sohocki MM, Sullivan LS, Mintz-Hittner HA, et al. A range of clinical phenotypes associated with mutations in CRX, a photoreceptor transcription-factor gene. AmJ HumGenet 1998;63:1307–1315.

11. Hull S, Arno G, Plagnol V, et al. The phenotypic variability of retinal dystrophies associated with mutations in CRX, with report of a novel macular dystrophy phenotype. Invest Ophthalmol Vis Sci 2014;55:6934–6944.

12. Griffith JF, DeBenedictis MJ, Traboulsi EI. A novel dominant CRX mutation causes adult-onset macular dystrophy. Ophthalmic Genet 2018;39:120–124.

13. Romdhane K, Vaclavik V, Schorderet DF, Munier FL, Viet Tran H. CRX-linked macular dystrophy with intrafamilial variable expressivity. Ophthalmic Genet 2018;1–5.

14. Stone EM. Leber congenital amaurosis - a model for efficient genetic testing of heterogeneous disorders: LXIV Edward Jackson Memorial Lecture. AmJ Ophthalmol 2007;144:791–811.

15. Koenekoop RK. An overview of Leber congenital amaurosis: a model to understand human retinal development. Surv Ophthalmol 2004;49:379–398.

16. Huang L, Xiao X, Li S, et al. CRX variants in cone-rod dystrophy and mutation overview. Biochem BiophRs Res Commun 2012;426:498–503.

17. Tran NM, Chen S. Mechanisms of blindness: Animal models provide insight into distinct CRX-associated retinopathies. Dev DRN 2014;243:1153–1166.

18. Swaroop A, Wang QL, Wu W, et al. Leber congenital amaurosis caused by a homozygous mutation (R90W) in the homeodomain of the retinal transcription factor CRX: direct evidence for the involvement of CRX in the development of photoreceptor function. Hum Mol Genet 1999;8:299–305.

19. Ibrahim MT, Alarcon-Martinez T, Lopez I, Fajardo N, Chiang J, Koenekoop RK. A complete, homozygous CRX deletion causing nullizygosity is a new genetic mechanism for Leber congenital amaurosis. Sci Rep 2018;8:5034.

20. Furukawa T, Morrow EM, Li T, Davis FC, Cepko CL. Retinopathy and attenuated circadian entrainment in Crx-deficient mice. Nat Genet 1999;23:466–470.

21. Ruzycki PA, Linne CD, Hennig AK, Chen S. Crx-L253X Mutation Produces Dominant Photoreceptor Defects in TVRM65 Mice. Invest Ophthalmol Vis Sci 2017;58:4644–4653.

22. Menotti-Raymond M, Deckman KH, David V, Myrkalo J, O’Brien SJ, Narfstrom K. Mutation discovered in a feline model of human congenital retinal blinding disease. Invest Ophthalmol Vis Sci 2010;51:2852–2859.

23. Occelli LM, Tran NM, Narfstrom K, Chen S, Petersen-Jones SM. CrxRdy Cat: A Large Animal Model for CRX-Associated Leber Congenital Amaurosis. Invest Ophthalmol Vis Sci 2016;57:3780–3792.

24. Hood DC, Lin CE, Lazow MA, Locke KG, Zhang X, Birch DG. Thickness of receptor and post-receptor retinal layers in patients with retinitis pigmentosa measured with frequency-domain optical coherence tomography. Invest Ophthalmol Vis Sci 2009;50:2328–2336.

25. Schneider CA, Rasband WS, Eliceiri KW. NIH Image to ImageJ: 25 years of image analysis. Nat Methods 2012;9:671–675.

26. RStudio Team (2015). RStudio: Integrated Development for R. RStudio I, Boston, MA URL http://www.rstudio.com/.

27. Staurenghi G, Sadda S, Chakravarthy U, Spaide RF, International Nomenclature for Optical Coherence Tomography P. Proposed lexicon for anatomic landmarks in normal posterior segment spectral-domain optical coherence tomography: the IN*OCT consensus. Ophthalmology 2014;121:1572–1578.

28. Ekstrom P, Sanyal S, Narfström K, Chader GJ, van VT. Accumulation of glial fibrillary acidic protein in Muller radial glia during retinal degeneration. Invest Ophthalmol Vis Sci 1988;29:1363–1371.

29. Sarthy PV, Fu M. Transcriptional activation of an intermediate filament protein gene in mice with retinal dystrophy. DNA 1989;8:437–446.

30. Linberg KA, Fariss RN, Heckenlively JR, Farber DB, Fisher SK. Morphological characterization of the retinal degeneration in three strains of mice carrying the rd-3 mutation. Vis Neurosci 2005;22:721–734.

31. Curtis R, Barnett KC, Leon A. An early-onset retinal dystrophy with dominant inheritance in the Abyssinian cat. Clinical and pathological findings. Invest OphthalmolVisSci 1987;28:131–139.

32. Chong NH, Alexander RA, Barnett KC, Bird AC, Luthert PJ. An immunohistochemical study of an autosomal dominant feline rod/cone dysplasia (Rdy cats). Exp Eye Res 1999;68:51–57.

33. Leon A, Curtis R. Autosomal dominant rod-cone dysplasia in the Rdy cat. 1. Light and electron microscopic findings. Exp Eye Res 1990;51:361–381.

34. Leon A, Hussain AA, Curtis R. Autosomal dominant rod-cone dysplasia in the Rdy cat. 2. Electrophysiological findings. Exp Eye Res 1991;53:489–502.

35. Nichols LL, 2nd, Alur RP, Boobalan E, et al. Two novel CRX mutant proteins causing autosomal dominant Leber congenital amaurosis interact differently with NRL. Hum Mutat 2010;31:E1472–1483.

36. Terrell D, Xie B, Workman M, et al. OTX2 and CRX rescue overlapping and photoreceptor-specific functions in the Drosophila eye. Dev Dyn 2012;241:215–228.

37. Roger JE, Hiriyanna A, Gotoh N, et al. OTX2 loss causes rod differentiation defect in CRX-associated congenital blindness. J Clin Invest 2014;124:631–643.

38. Yeh CY, Koehl KL, Harman CD, et al. Assessment of Rod, Cone, and Intrinsically Photosensitive Retinal Ganglion Cell Contributions to the Canine Chromatic Pupillary Response. Invest Ophthalmol Vis Sci 2017;58:65–78.

39. Hippert C, Graca AB, Barber AC, et al. Muller glia activation in response to inherited retinal degeneration is highly varied and disease-specific. PloS one 2015;10:e0120415.

40. Hollingsworth TJ, Gross AK. Defective trafficking of rhodopsin and its role in retinal degenerations. Int Rev Cell Mol Biol 2012;293:1–44.

41. Gao J, Cheon K, Nusinowitz S, et al. Progressive photoreceptor degeneration, outer segment dysplasia, and rhodopsin mislocalization in mice with targeted disruption of the retinitis pigmentosa-1 (Rp1) gene. Proc Natl Acad Sci U S A 2002;99:5698–5703.

42. Hollingsworth TJ, Gross AK. The severe autosomal dominant retinitis pigmentosa rhodopsin mutant Ter349Glu mislocalizes and induces rapid rod cell death. J Biol Chem 2013;288:29047–29055.

43. Bandyopadhyay M, Kono M, Rohrer B. Explant cultures of Rpe65-/-mouse retina: a model to investigate cone opsin trafficking. Mol Vis 2013;19:1149–1157.

44. Rana T, Shinde VM, Starr CR, et al. An activated unfolded protein response promotes retinal degeneration and triggers an inflammatory response in the mouse retina. Cell Death Dis 2014;5:e1578.

45. Athanasiou D, Aguila M, Bevilacqua D, Novoselov SS, Parfitt DA, Cheetham ME. The cell stress machinery and retinal degeneration. FEBS letters 2013;587:2008–2017.

46. McMullen RJ, Jr., Davidson MG, Gilger BC. The effect of 1 % tropicamide-induced mydriasis and cycloplegia on spherical refraction of the adult horse. Vet Ophthalmol 2014;17:120–125.

47. Twelker JD, Mutti DO. Retinoscopy in infants using a near noncycloplegic technique, cycloplegia with tropicamide 1%, and cycloplegia with cyclopentolate 1%. Optom Vis Sci 2001;78:215–222.

48. Schaeffel F, Feldkaemper M. Animal models in myopia research. Clin Exp Optom 2015;98:507–517.

49. Konrade KA, Hoffman AR, Ramey KL, Goldenberg RB, Lehenbauer TW. Refractive states of eyes and associations between ametropia and age, breed, and axial globe length in domestic cats. Am J Vet Res 2012;73:279–284.

50. Whitmore WG. Congenital and developmental myopia. Eye (Lond) 1992;6 (Pt 4):361–365.

51. Ritchey ER, Zelinka C, Tang J, et al. Vision-guided ocular growth in a mutant chicken model with diminished visual acuity. Exp Eye Res 2012;102:59–69.

52. Rada JA, Shelton S, Norton TT. The sclera and myopia. Exp Eye Res 2006;82:185–200.

