## Supplementary materials for "Cat LCA-*CRX* model, homozygous for an antimorphic mutation has a unique phenotype"

**Occelli et al**

**Supplementary Figure S1.**

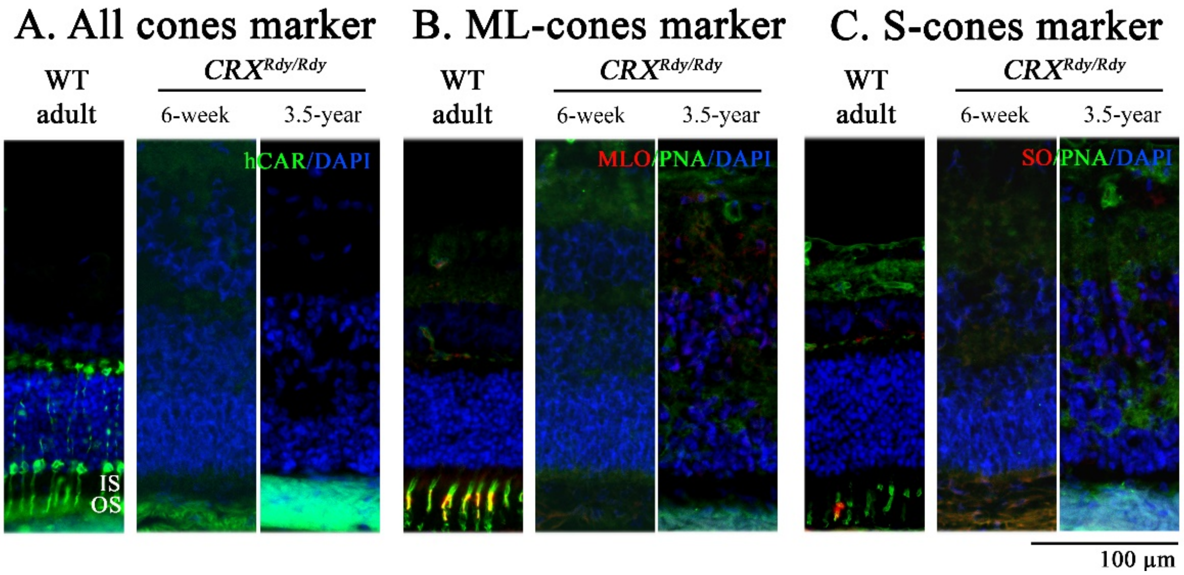

**Supplemental Figure S1.** Immunohistochemistry for cone markers (human cone arrestin, ML opsin, S-opsin and the lectin PNA). There was no labeling in the *CRX*$^{Rdy/Rdy}$ cat retina. WT control retina shown for comparison. hCAR labels the entire cone photoreceptor cell body in WT cats. PNA labels the cone matrix in WT cats. MLO and SO are visible within the outer segment of the cone photoreceptor cells in WT cats.

### Supplementary Figure S2.

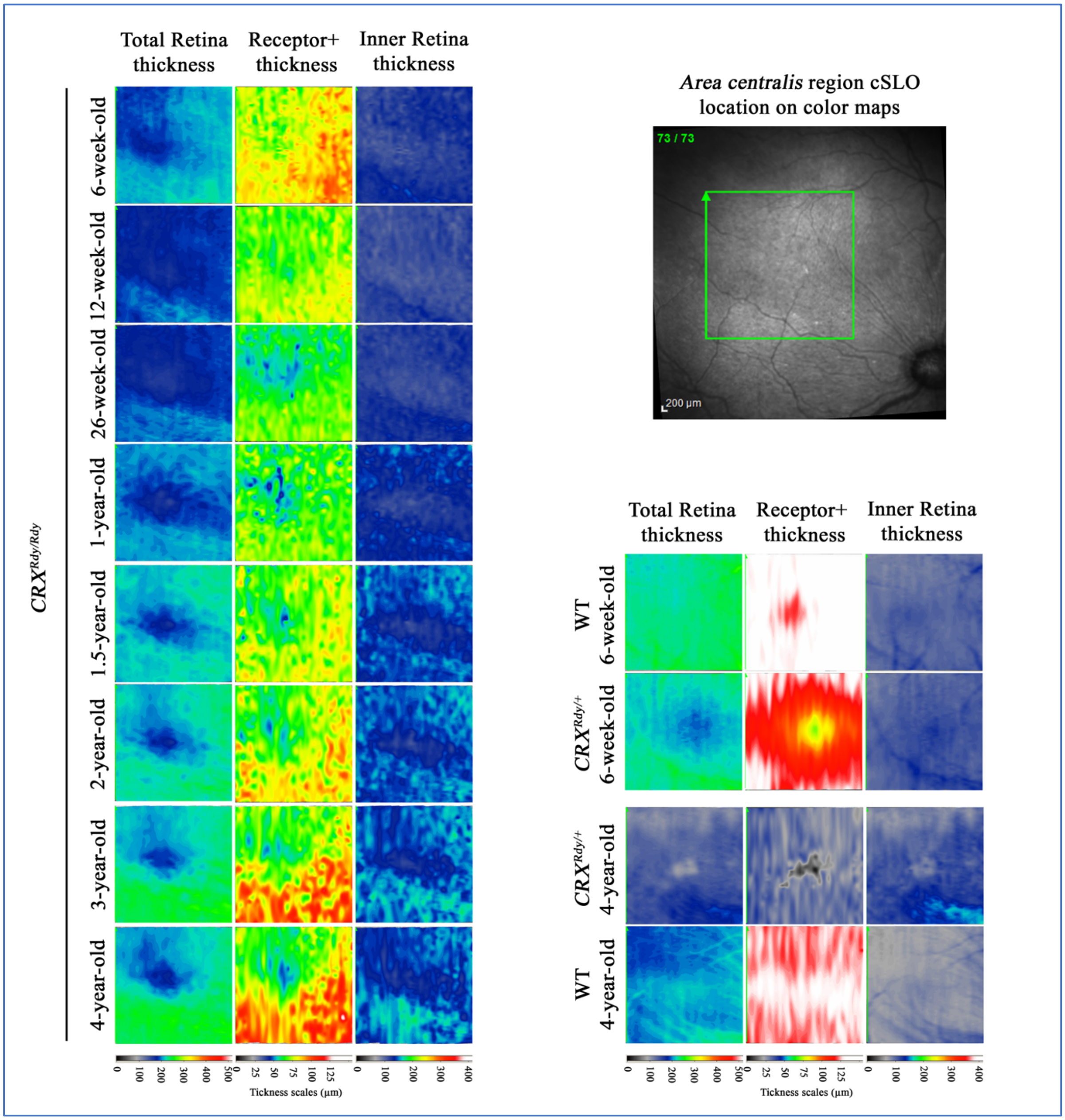

**Supplemental Figure S2.** Total retinal (TR), Receptor+ (REC+) and Inner retina (IR) thicknesses color map in the *area centralis*. The left panel shows color (heat) maps for a representative $CRX^{Rdy/Rdy}$ cat from 6 weeks to 4 years of age. The right top panel shows the retinal location of the color map. The bottom right panel shows the color map for representative $CRX^{Rdy/+}$and WT cats at 6 weeks and 4 years of age. Note the thinner TR and REC+ in the $CRX^{Rdy/Rdy}$ cat from an early age compared to WT cats. The IR is thicker than that of the $CRX^{Rdy/+}$ and WT cats. With disease progression, thickening surrounding the center of the *area centralis* can be seen in the $CRX^{Rdy/Rdy}$ cat, leading to slight TR and REC+ thickening, which is in contrast with the severe thinning of retinal layers in the $CRX^{Rdy/+}$ cats at 4 year of age.

**Supplementary Table S1.**

| *CRX*$^{Rdy/Rdy}$ | | | WT | | | *CRX*$^{Rdy/+}$ | | |
|---|---|---|---|---|---|---|---|---|
| **Age (years)** | **GL both eyes** | **C-AC, L, PS both eyes** | **Age (years)** | **GL both eyes** | **C-AC, L, PS both eyes** | **Age (years)** | **GL both eyes** | **C-AC, L, PS both eyes** |
| 0.074 | 1 | | 0.071 | 3 | | 0.038 | 1 | |
| 0.082 | 1 | | 0.074 | 1 | | 0.071 | 2 | |
| 0.112 | 3 | 3 | 0.09 | 3 | | 0.074 | 2 | |
| 0.118 | 2 | | 0.109 | 3 | | 0.09 | 2 | |
| 0.148 | 1 | | 0.112 | 1 | 1 | 0.109 | 3 | |
| 0.165 | 1 | | 0.118 | 2 | | 0.112 | 1 | 1 |
| 0.192 | 1 | 1 | 0.123 | 2 | | 0.118 | 2 | |
| 0.2 | 1 | 1 | 0.129 | 2 | | 0.129 | 1 | |
| 0.227 | 1 | 1 | 0.137 | 3 | | 0.137 | 2 | |
| 0.23 | 2 | 1 | 0.14 | 1 | | 0.165 | 1 | |
| 0.274 | 1 | 1 | 0.142 | 1 | | 0.175 | 2 | |
| 0.288 | 2 | 1 | 0.148 | 1 | | 0.194 | 1 | |
| 0.297 | 2 | | 0.165 | 1 | | 0.197 | 2 | |
| 0.337 | 2 | 1 | 0.175 | 3 | | 0.2 | 1 | |
| 0.384 | 2 | 1 | 0.192 | 1 | | 0.225 | 2 | 1 |
| 0.412 | 1 | | 0.194 | 1 | | 0.227 | 1 | 1 |
| 0.474 | 1 | | 0.197 | 1 | | 0.233 | 1 | |
| 0.499 | 2 | 2 | 0.2 | 1 | | 0.236 | 1 | |
| 0.567 | 1 | | 0.203 | 1 | | 0.238 | 1 | |
| 0.778 | 1 | | 0.225 | 1 | | 0.274 | 1 | |
| 1 | 2 | 1 | 0.227 | 2 | 1 | 0.288 | 4 | |
| 1.5 | 1 | 1 | 0.233 | 1 | | 0.297 | 2 | |
| 2 | 2 | 2 | 0.236 | 1 | | 0.329 | 1 | |
| | | | 0.238 | 2 | | 0.337 | 1 | |
| | | | 0.271 | 1 | | 0.377 | 3 | 1 |
| | | | 0.274 | 1 | | 0.384 | 3,5 | |
| | | | 0.288 | 3 | | 0.412 | 2 | |
| | | | 0.297 | 2 | | 0.499 | 3 | |
| | | | 0.329 | 2 | | 0.567 | 1 | |
| | | | 0.337 | 2 | | 0.775 | 1 | |

| | | | | | | | | |
|---|---|---|---|---|---|---|---|---|
| | | | 0.377 | 3 | | 1 | 4 | 1 |
| | | | 0.384 | 1 | | 1.5 | 1 | 1 |
| | | | 0.412 | 2 | | 2.5 | 1 | 1 |
| | | | 0.479 | 1 | | 3 | 6 | 1 |
| | | | 0.499 | 1 | 1 | 4 | 2 | 2 |
| | | | 0.567 | 2 | | 4.5 | 2 | |
| | | | 0.775 | 1 | | 4.75 | 1 | 1 |
| | | | 1 | 9 | 5 | 5 | 2 | |
| | | | 1.5 | 2 | 1 | 9 | 2 | |
| | | | 2 | 1 | 1 | 10.1 | 2 | 2 |
| | | | 6 | 1 | | | | |
| **Total different animals** | | | | | | | | |
| | 6 | 5 | | 24 | 9 | | 30 | 12 |

**Supplementary Table S1. Experiments and numbers of animal for globe length in $CRX^{Rdy/Rdy}$, $CRX^{Rdy/+}$and WT cats**

**GL both eyes:** Axial globe length measurements in both eyes of one individual.
**C-AC, L, PS both eyes:** Cornea-Anterior chamber width, Lens width and Posterior Segment in both eyes of one individual.

**Supplementary Table S2.**

**IOP**

| **Age (years)** | ***CRX*** $^{Rdy/Rdy}$ | **WT** |
|---|---|---|
| 0.6 | | 2 |
| 0.8 | 2 | |
| 0.9 | | 1 |
| 2.5 | 1 | |
| 4.1 | 1 | 1 |
| 6.2 | 1 | |
| **Total different animals** | **6** | **5** |

**Refraction**

| **Age (years)** | ***CRX*** $^{Rdy/Rdy}$ | **WT** | ***CRX*** $^{Rdy/+}$ |
|---|---|---|---|
| 0.6 | | 2 | |
| 0.8 | 2 | | 1 |
| 0.9 | | 1 | |
| 1.4 | | | 1 |
| 2.5 | 2 | | |
| 3.2 | | 1 | |
| 3.7 | | | 1 |
| 4.1 | 1 | 1 | |
| 4.6 | | | 1 |
| 6.2 | 1 | | |
| 6.6 | | | 1 |
| 6.7 | | | 1 |
| **Total different animals** | **6** | **5** | **6** |

**qRT-PCR**

| **Age (weeks)** | ***CRX*** $^{Rdy/Rdy}$ | **WT** | ***CRX*** $^{Rdy/+}$ |
|---|---|---|---|
| 2 | 3 | 3 | 3 |

**Western**

| **Age (weeks)** | ***CRX*** $^{Rdy/Rdy}$ | **WT** |
|---|---|---|
| 2 | 2 | 3 |

| | | **Immunohistochemistry** | **Histology** |
|---|---|---|---|
| **Age (years)** | **Age (weeks)** | ***CRX*** $^{Rdy/Rdy}$ | ***CRX*** $^{Rdy/Rdy}$ |
| 0.04 | 2 | 2 | 2 |
| 0.12 | 6 | 2 | 2 |
| 0.23 | 12 | 2 | 2 |
| 0.38 | 20 | 3 | 3 |
| 2 | | 1 | 1 |

| | | | |
|---|---|---|---|
| 3.5 | | 1 | 1 |

**Supplementary Table S2. Experiments and numbers of animal for IOP, refraction, qRT-PCR, Western blot, immunohistochemistry and histology in *CRX*$^{Rdy/Rdy}$, *CRX*$^{Rdy/+}$ and WT cats**

**Supplementary Table S3.**

| Age (years) | Age (weeks) | Fluorescein angiography | ERG | SD-OCT |
|---|---|---|---|---|
| 0.08 | 4 | | 2 | 2 |
| 0.12 | 6 | | 6 | 4 |
| 0.15 | 8 | | 2 | 2 |
| 0.19 | 10 | | 4 | 3 |
| 0.23 | 12 | | 5 | 8 |
| 0.29 | 15 | | 4 | 4 |
| 0.34 | 17.5 | | 1 | |
| 0.38 | 20 | | 7 | 6 |
| 0.5 | 26 | 2 | | 3 |
| 0.58 | | | | 1 |
| 0.77 | | | | 1 |
| 1 | | 1 | | 4 |
| 1.5 | | | | 3 |
| 2 | | 1 | | 3 |
| 3 | | 1 | | 2 |
| 3.5 | | | | 1 |
| 4 | | 1 | | 2 |
| 5 | | | | 1 |
| **Total different animals** | | **4** | **10** | **12** |

**Supplementary Table S3. Experiments and numbers of animal for fluorescein angiography, ERG and SD-OCT in *CRX*$^{Rdy/Rdy}$ cats**

### Supplementary Table S4.

| Antibody – Source | Type | Primary Dilution | Secondary Antibody – Source | Secondary Dilution |
|---|---|---|---|---|
| **Hcar** (Human cone arrestin) Dr. Cheryl Craft; LUMIJ, University of Southern California, Los Angeles, CA, USA | Polyclonal rabbit | 1:10,000 | Alexa Fluor 488 Goat anti-rabbit IgG Life technologies, Carlsbad, CA, USA | 1:500 |
| **PNA** (Biotinylated Peanut Agglutinin) Vector Labs Inc., Burlin-game, CA, USA | Biotinylated Lectin | 1:500 | Alexa Fluor 488 Streptavadin Life technologies, Carlsbad, CA, USA | 1:500 |
| **ML-opsin** (Anti-Opsin, Red/Green; Medium/ Long wavelength cone opsin) Millipore Corp., Billerica, MA, USA | Polyclonal rabbit | 1:1,000 | Alexa Fluor 568 or 594 Goat anti-rabbit IgG Life technologies, Carlsbad, CA, USA | 1:500 |
| **S-opsin** (Anti-Opsin, Blue; Short wavelength cone opsin) Millipore Corp., Billerica, MA, USA | Polyclonal rabbit | 1:1,000 | Alexa Fluor 568 or 594 Goat anti-rabbit IgG Life technologies, Carlsbad, CA, USA | 1:500 |
| **RetP1** (Rhodopsin Ab-1) Thermo Scientific, Rockford, IL, USA | Monoclonal mouse | 1:2 | Alexa Fluor 594 Goat anti-mouse IgG Life technologies, Carlsbad, CA, USA | 1:500 |
| **GFAP** (Anti-Glial Fibrillary Acidic Protein) Cell Signaling Technology Inc., Danvers, MA, USA | Monoclonal mouse | 1:300 | Alexa Fluor 594 Rabbit anti-mouse IgG Life technologies, Carlsbad, CA, USA | 1:500 |
| **PKCα** (Protein Kinase C-alpha) BD Biosciences, San Jose, CA, USA | Monoclonal mouse | 1:500 | Alexa Fluor 594 Goat anti-mouse IgG Life technologies, Carlsbad, CA, USA | 1:500 |
| **NeuN** (Neuron-Specific Nuclear Protein) Millipore Corp., Billerica, MA, USA | Monoclonal mouse | 1:2,000 | Alexa Fluor 488 Goat anti-mouse IgG Life technologies, Carlsbad, CA, USA | 1:500 |
| **CRX** (Cone rod homebox) Sigma-Aldrich, St Louis, MO, USA | Monoclonal mouse | 1:20,000 | Alexa Fluor 568 Goat anti-mouse IgG Life technologies, Carlsbad, CA, USA | 1:500 |

**Supplementary Table S4. List of antibodies used for IHC – their origins and dilutions**

### Supplementary Table S5.

| Primer name | Forward primer | Reverse primer | Amplicon size (bp) |
|---|---|---|---|
| *ARR3* | 5' CGTTGTCCTGTATTCCCTAGAC 3' | 5' GCTAGAGGCCAGATTAGTATCAC 3' | 190 |
| *RHO* | 5' GGTGCCCTACGCCAGCGTG 3' | 5' CAGTGGGTTCTTGCCACAG 3' | 190 |
| *MOP* | 5' TGTCTCCTTGTGTGGGATCA 3' | 5' GTACGAGCTGCCACTGAACA 3' | 257 |
| *SO* | 5' AGTCAGCCTCAACCCAGAA 3' | 5' CACCATCTCCATGATGCAAG 3' | 326 |
| *CRX Total* | 5' AAGACTCAGTACCCGGATGTGTA 3' | 5' GGGGCTGTAGGAGTCTGAGAT 3' | 223 |
| *OTX2* | 5' GCTAGACGTGCTGGAAGCTC 3' | 5' GGGCTGGAGAGGTCTTCTTT 3' | 209 |
| *NRL* | 5' CCCACAGCTACTACCCAGGA 3' | 5' TCACACCACTCCCTCTCCTC 3' | 207 |
| *NR2E3* | 5' CATCCCCATACTCCTCCTCA 3' | 5' GAGGCAGGGACCACTGTATG 3' | 198 |
| *TRB2* | 5' GACTCCGAACTTGTCGCATT 3' | 5' TTTGGTTGATGTTGCTGTGG 3' | 197 |
| *RORB* | 5' GAGGAATGCAGATGTTCAAGG 3' | 5' GCTTCTGGACCTTCCTTGGT 3' | 177 |
| *TUBA1B* | 5' GCTCTATTGCCTGGAACACG 3' | 5' CATCTTCCTTGCCCGTGATG 3' | 230 |
| *GAPDH* | 5' GGTCTTCACCACCATGGAGA 3' | 5' TGGACTGTGGTCATGAGTCC 3' | 237 |

**Supplementary Table S5. Primer sequences for qRT-PCR assays**

### Supplementary Table S6.

| mRNA | ***Crx***$^{Rdy/Rdy}$ ***vs Crx***$^{Rdy/+}$ | ***Crx***$^{Rdy/Rdy}$ ***vs*** **WT** | ***Crx***$^{Rdy/+}$ ***vs*** **WT** |
|---|---|---|---|
| ***Arr3*** | 27.4x less (P = 0.003) | 1724.1x less (P = 0.004) | 62.9x less (P = 0.002) |
| ***SO*** | 15.8x less (P = 0.010) | 62.7x less (P = 0.004) | 4.0x less (P = 0.026) |
| ***MOP*** | Not detectable in *Crx*$^{Rdy/Rdy}$ (P = 0.001) | Not detectable in *Crx*$^{Rdy/Rdy}$ (P ≤ 0.001) | 4.3x less (P = 0.002) |
| ***Rho*** | 10.9x less (P ≤ 0.001) | 32.5x less (P = 0.004) | 3.0x less (P ≤ 0.001) |
| **total *Crx*** | No significant difference (P = 0.792) | 1.8x more (P = 0.032) | 1.8x more (P = 0.002) |
| ***Otx2*** | No significant difference (P = 0.119) | 2.4x more (P ≤ 0.001) | 1.9x more (P ≤ 0.001) |
| ***Nrl*** | 2.3x less (P = 0.029) | No significant difference (P = 0.058) | No significant difference (P = 0.603) |
| ***Nr2e3*** | 2.1x less (P ≤ 0.001) | 1.9x less (P ≤ 0.001) | No significant difference (P = 0.295) |
| ***Trβ2*** | No significant difference (P = 0.507) | 1.4x more (P = 0.030) | No significant difference (P = 0.069) |
| ***Rorβ*** | 1.3x more (P = 0.021) | 1.3x more (P = 0.049) | No significant difference (P = 0.822) |

**Supplementary Table S6. *P* values for qRT-PCR assays results**
